# Synaptic alterations in pyramidal cells following genetic manipulation of neuronal excitability in monkey prefrontal cortex

**DOI:** 10.1101/2024.06.12.598658

**Authors:** Guillermo Gonzalez Burgos, Takeaki Miyamae, Yosuke Nishihata, Olga L Krimer, Kirsten Wade, Kenneth N Fish, Dominique Arion, Zhao-Lin Cai, Mingshan Xue, William R Stauffer, David A Lewis

## Abstract

In schizophrenia, layer 3 pyramidal neurons (L3PNs) in the dorsolateral prefrontal cortex (DLPFC) are thought to receive fewer excitatory synaptic inputs and to have lower expression levels of activity-dependent genes and of genes involved in mitochondrial energy production. In concert, these findings from previous studies suggest that DLPFC L3PNs are hypoactive in schizophrenia, disrupting the patterns of activity that are crucial for working memory, which is impaired in the illness. However, whether lower PN activity produces alterations in inhibitory and/or excitatory synaptic strength has not been tested in the primate DLPFC. Here, we decreased PN excitability in rhesus monkey DLPFC *in vivo* using adeno-associated viral vectors (AAVs) to produce Cre recombinase-mediated overexpression of Kir2.1 channels, a genetic silencing tool that efficiently decreases neuronal excitability. In acute slices prepared from DLPFC 7-12 weeks post-AAV microinjections, Kir2.1-overexpressing PNs had a significantly reduced excitability largely attributable to highly specific effects of the AAV-encoded Kir2.1 channels. Moreover, recordings of synaptic currents showed that Kir2.1-overexpressing DLPFC PNs had reduced strength of excitatory synapses whereas inhibitory synaptic inputs were not affected. The decrease in excitatory synaptic strength was not associated with changes in dendritic spine number, suggesting that excitatory synapse quantity was unaltered in Kir2.1-overexpressing DLPFC PNs. These findings suggest that, in schizophrenia, the excitatory synapses on hypoactive L3PNs are weaker and thus might represent a substrate for novel therapeutic interventions.

**Significance Statement:** In schizophrenia, dorsolateral prefrontal cortex (DLPFC) pyramidal neurons (PNs) have both transcriptional and structural alterations that suggest they are hypoactive. PN hypoactivity is thought to produce synaptic alterations in schizophrenia, however the effects of lower neuronal activity on synaptic function in primate DLPFC have not been examined. Here, we used, for the first time in primate neocortex, adeno-associated viral vectors (AAVs) to reduce PN excitability with Kir2.1 channel overexpression and tested if this manipulation altered the strength of synaptic inputs onto the Kir2.1-overexpressing PNs. Recordings in DLPFC slices showed that Kir2.1 overexpression depressed excitatory (but not inhibitory), synaptic currents, suggesting that, in schizophrenia, the hypoactivity of PNs might be exacerbated by reduced strength of the excitatory synapses they receive.

## Introduction

In schizophrenia, layer 3 pyramidal neurons (L3PNs) in the dorsolateral prefrontal cortex (DLPFC) have a lower density of dendritic spines (Garey et al., 1998; Glantz and Lewis, 2000), the main sites of excitatory synaptic inputs onto PNs. These neurons also express lower levels of activity-dependent genes (Weickert et al., 2003; Hashimoto et al., 2005) and of genes involved in mitochondrial energy production (Arion et al., 2015; Arion et al., 2017; Glausier et al., 2020). In concert, these morphological and molecular alterations suggest that schizophrenia is associated with lower activity of DLPFC L3PNs (Lewis et al., 2012). Lower DLPFC neuron activity is consistent with data from fMRI (Minzenberg et al., 2009; Smucny et al., 2023) and EEG studies (Cho et al., 2006; Boudewyn et al., 2020) showing that DLPFC activation or gamma oscillation power during cognitive tasks are reduced in schizophrenia.

Lower PN firing might affect the activity-dependent regulation of synaptic strength (Wu et al., 2022; Debanne and Inglebert, 2023), with both compensatory and deleterious effects. For instance, a compensatory decrease in inhibitory synaptic strength might maintain excitation and inhibition in balance, but also disrupt the inhibition-based network oscillations (Gonzalez-Burgos et al., 2015a) associated with working memory (Miller et al., 2018) which is impaired in schizophrenia (Smucny et al., 2022). Lower PN activity could also affect the mechanisms regulating glutamate synapse strength (Wu et al., 2022; Debanne and Inglebert, 2023), further disrupting DLPFC microcircuit function. Although crucial to understand the mechanisms of microcircuit dysfunction in schizophrenia, the effects of reduced PN activity on inhibitory and/or excitatory synaptic inputs to these neurons has not been tested in primate DLPFC.

To address this question, we decreased PN excitability chronically in area 46 of rhesus monkey DLPFC *in vivo* using adeno-associated viral vector (AAV)-mediated overexpression of Kir2.1 channels, a genetic silencing tool (Wiegert et al., 2017) that efficiently decreased the excitability, and thus activity, of cortical neurons in Cre mouse lines (Xue et al., 2014). Rodents, however, do not possess a neocortical area analogous or homologous to the expanded primate DLPFC (Ma et al., 2022; Preuss and Wise, 2022). Moreover, during working memory tasks, primate DLPFC neurons display activity patterns of exceptional complexity and duration not observed in other cortical areas or species (Constantinidis et al., 2018; Lundqvist et al., 2018). Therefore, reducing neuronal excitability *in vivo* may affect synaptic communication in the primate DLPFC via unique effects that are not reproducible using rodents as model.

To assess the effects of *in vivo* Kir2.1 overexpression we employed patch clamp recordings from PNs in acute slices prepared 7-12 weeks post-AAV microinjections. We found that Kir2.1 overexpression largely reduced PN excitability via highly specific effects of the Kir2.1 channels encoded by the AAVs. Furthermore, recordings of synaptic currents showed that Kir2.1 overexpression reduced the strength of excitatory but not inhibitory synaptic inputs on DLPFC PNs. Additionally, in the Kir2.1-overexpressing DLPFC PNs with decreased excitatory synaptic strength, dendritic spine number was not changed, showing that the alterations in excitatory synapse function were not accompanied by changes in excitatory synapse quantity. These findings suggest that, in schizophrenia, lower PN activity is not sufficient to cause persistent changes in inhibitory synapse strength but might affect the strength of the excitatory synapses in DLPFC PNs.

## Materials and Methods

### Plasmids and AAV vector preparation

The plasmid pAAV-EFla-DIO-Myc-Kir2.1 E224G Y242F-P2A-dTomato-WPRE-bGHpA was cloned and used to produce AAV9 vectors at UPENN vector core. We purchased AAV2-CamKII-EGFP-T2A-iCre from Vector Biolabs (VB1918). Both vectors were stored at -80 C. Preliminary studies with AAV2-CamKII-EGFP-T2A-iCre examined expression levels after injection titers 10^11^, 10^12^, and 10^13^. Based on these results, we selected 10^12^ as the titer that resulted in good expression mostly in pyramidal neurons. Directly prior to surgery, we combined AAV2-CamKII-iCre-GFP-WPRE, AAV9-EFla-DIO-Myc-Kir2.1 E224G Y242F-P2A-dTomato-WPRE-bGHpA, and sterile saline such that the final titers were 10^12^ and 10^13^, respectively. Using these titers, the spatial distribution of AAV-mediated expression, assessed via the dTomato fluorescence (Extended Data Fig 1), was very similar to that observed in previous studies using the same AAV serotypes to drive gene expression in the macaque monkey neocortex (Inoue et al., 2015; Watakabe et al., 2015).

### Animals and surgery

All animal procedures were performed in accordance with the National Institutes of Health Guide for the Care and Use of Laboratory Animals and approved by the University of Pittsburgh’s Institutional Animal Care and Use Committee (IACUC) (Protocol ID, 19024431). Four male rhesus monkeys aged 2.5-4 years old were single- or pair-housed with a 12h-12h light-dark cycle. AAV injections were done under general anesthesia. In Monkeys B, D, and C, we created a cranial window that enabled us to visualize the principal sulcus (PS) and arcuate sulcus (AS). In each injections track, we injected 5-20 μl of AAV solutions. Injections were made in both the dorsal and ventral banks of the PS. In monkey H, we used MRI guided robotic surgery to inject 15 μl per track in six tracks, all parallel to the PS, located in the dorsal and ventral banks. Monkeys B and D were assigned to experiments studying inhibitory synaptic currents, whereas monkeys C and H, were allocated to experiments studying excitatory synaptic currents.

### Multiplex Fluorescent RNA in situ Hybridization (FISH)

A fresh-frozen coronal tissue section (16 micrometer-thick) containing left DLPFC area 46 was obtained from one of the rhesus monkeys around AAV injections placed in the left hemisphere, whereas DLPFC tissue from the right hemisphere of the same animal was used for electrophysiology experiments. The tissue section was thaw-mounted in a cryostat, onto SuperFrost slides (ThermoFisher), and stored at –80°C. We performed multiplex FISH, using probes designed by Advanced Cell Diagnostics, Inc (ACDbio; Hayward, CA, USA) to detect mRNAs encoding vesicular glutamate transporter 1, (VGluT1; *SLC17A7* gene; Rhesus macaque probe #424831) and Kir2.1 (*KCNJ2* gene; Mus musculus probe #476261).

A commercially available kit FISH assay from ACDbio was used to detect both mRNA targets. Briefly, after removal from –80°C storage, a slide-mounted section was immediately immersed in ice-cold 4% paraformaldehyde for 12 minutes, dehydrated through a series of ethanol washes (50%, 70%, 100%, and repeated 100% ethanol for a few seconds each), followed by pretreatment with protease Plus (ACDbio) for 12 minutes at room temperature. The section was then incubated with both probes diluted 1:50 in blank probe diluent for 1 hours at 40°C. Amplification and labeling steps were conducted according to the manufacturer’s instructions. Fluorophore Opal 570 (Akoya Biosciences) was assigned to KCNJ2 mRNA and Opal 520 was assigned to VGluT1 mRNA. Sections were then quickly dehydrated in 100% ethanol and inspected under the microscope to detect VGluT1 and KCNJ2 labeling.

### Electrophysiology studies

#### Solutions employed

All the reagents used to prepare solutions were purchased from Sigma-Aldrich (www.sigmaaldrich.com). The concentrations below are in mM units.

#### Sucrose-ACSF

Sucrose 200, NaCl 15, KCl 1.9, Na_2_HPO_4_ 1.2, NaHCO_3_ 33, MgCl_2_ 6, CaCl_2_ 0.5, glucose 10 and kynurenic acid 2; pH 7.3–7.4 when bubbled with 95% O_2_-5% CO_2_.

#### ACSF

NaCl 125, KCl 2.5, Na_2_HPO_4_ 1.25, glucose 10, NaHCO_3_ 25, MgCl_2_ 1 and CaCl_2_ 2, pH 7.3–7.4 when bubbled with 95% O_2_-5% CO_2_.

#### High intracellular chloride pipette solution

KGluconate 60; KCl 70; EGTA 0.2; HEPES 10; MgATP 4; NaGTP 0.3, NaPhosphocreatine 14, biocytin 0.4 %, pH 7.2-7.3, adjusted with KOH.

#### Standard intracellular chloride pipette solution

KGluconate 120; KCl 10; EGTA 0.2; HEPES 10; MgATP 4; NaGTP 0.3, NaPhosphocreatine 14, biocytin 0.4 %, pH 7.2-7.3, adjusted with KOH.

### Acute brain slice preparation

After allowing 7-12 weeks for AAV-mediated expression, tissue blocks containing the AAV injection sites in the principal sulcus region of DLPFC area 46 (right hemisphere) were obtained after the animals were deeply anesthetized and perfused transcardially (Gonzalez-Burgos et al., 2015b) with ice-cold sucrose-modified artificial cerebro-spinal fluid (sucrose-ACSF). The location of the AAV injection sites in the tissue blocks used for slice preparation was, first, estimated using images obtained during the microinjection surgeries, and, second, verified by visualization using a Dual Fluorescent Protein Flashlight (Nightsea, Hatfield PA) which allowed detecting red fluorescent spots at the surface of the tissue blocks for some, albeit not all, the injection sites.

Three hundred µm thick slices were cut in the coronal plane starting from the rostral end of the tissue blocks using a vibrating microtome (VT1200S, Leica Microsystems) while submerged in ice-cold sucrose-ACSF. Immediately after cutting, the slices were transferred to an incubation chamber filled with room temperature ACSF. Electrophysiological recordings were obtained 1-14 hours after tissue slicing was completed.

#### Targeting of cells for recordings in the acute brain slices

Slices were placed in recording chambers superfused at 2-3 ml/min with oxygenated ACSF at 30-32 °C. To record sIPSCs or sEPSCs, the AMPAR or GABA_A_R antagonists CNQX (10 µM) or SR95531 (10 µM) were added to the ACSF, respectively. Whole-cell recordings were obtained from PNs identified visually by infrared differential interference contrast (IR-DIC) and epifluorescence video microscopy using Olympus or Zeiss microscopes equipped with CCD video cameras (EXi Aqua, Q-Imaging). Cells identified as PNs with IR-DIC were visualized with epifluorescence microscopy to detect the red fluorescence from the dTomato encoded by the Flox-Kir2.1-dTomato vector. PNs with (dT+) or without (dT-) detectable red fluorescence, were targeted for recording to determine the effects of Kir2.1 overexpression. Cells were targeted for recordings in layers 3 to 5/6 of either medial or lateral banks of the principal sulcus in DLPFC area 46, depending on the anatomical location of the injection sites and on the laminar distribution of the dT+ PNs in each slice. Slices apparently containing an AAV injection site typically had substantial numbers of dT+ neurons but also showed evidence of damage by the placement of the injection needles and were not used in our experiments. Cells were usually targeted for recording starting with adjacent slices immediately rostral or caudal to a slice apparently containing an injection site. Whereas the number of dT+ neurons clearly declined as a function of distance from an injection site, the dTomato fluorescence signal intensity in single cells did not consistently vary with distance from an injection site. Moreover, we could not distinguish an obvious relation between the magnitude of the biophysical effects of Kir2.1 overexpression and the location of the recorded cells relative to an injection. To the contrary, the size of the Kir2.1 overexpression effect was usually markedly different in nearby dT+ PNs recorded in a single slice and the same cortical layer, as reflected by, for instance, highly different rheobase values. These observations suggest that the magnitude of the Kir2.1 overexpression effect in single cells was determined by variable numbers of AAV particles infecting single PNs, cell-to-cell variability in the efficiency of transcription, or both, but not by the overall spatial location of the neurons relative to an injection site. Cre recombinase activity was less likely to be a determinant factor, because we microinjected an excess of AAV2-CamKII-EGFP-T2A-iCre vector relative to AAV9-EFla-DIO-Myc-Kir2.1 E224G Y242F-P2A-dTomato-WPRE-bGHpACAMK2A-Cre-GFP vector particles.

#### Electrophysiological recordings and analysis

Recording pipettes had 3-5 MΩ resistance when filled with either of the two different solutions. To record sIPSCs or sEPSCs, pipettes were filled with high intracellular chloride or standard intracellular chloride KGluconate-based pipette solutions, respectively. Biocytin (0.4 %) was included to fill the cells during recordings and later stain the tissue with the DAB method (González-Burgos et al., 2019).

#### Current clamp

Recordings and data analysis were conducted with Multiclamp 700A or 700B amplifiers (Axon Instruments) operating in bridge mode with pipette capacitance neutralization (Miyamae et al., 2017; González-Burgos et al., 2019). Recordings were included in data analysis only if the resting membrane potential was ≤-60 mV. Membrane properties were measured using families of 0.5 or 1 s current steps, starting from -110 pA until reaching at least 250 pA above rheobase, incrementing by 10 pA (2 repeats per current level). The procedures used to measure membrane properties (Miyamae et al., 2017; González-Burgos et al., 2019) are summarized below.

#### Dynamic clamp

Most of the dynamic clamp experiments were performed in PNs from mouse medial PFC used as a model, whereas dynamic clamp experiments in dT-PNs from DLPFC area 46 (n=2) revealed very similar effects. Slices from mouse PFC were prepared as described elsewhere (Miyamae et al., 2017). Dynamic clamp was implemented with the software-based system employed in the StdpC dynamic clamp system (Nowotny et al., 2006; Kemenes et al., 2011) and available using Signal 7 and Power1401-3A data acquisition interfaces (Cambridge Electronic Design, Cambridge, UK). Kir conductance (G*_Kir_*) was simulated using Hodgkin-Huxley (Tau) or Hodgkin-Huxley (Alpha/Beta) models with G*_Kir,max_* values of 1, 5, 10 or 20 nS, reversal potential E_r_=-80 mV. As in previous studies (Amarillo et al., 2018; Szegedi et al., 2023), the G*_Kir_* models used in dynamic clamp did not include an inactivation gating variable (h). The parameters for the activation variable (m) were, in the Tau model (Szegedi et al., 2023): p=1, V_th_=-28 mV, V_slope_=-7.5 mV, Tau_0_=2 ms, Tau_a_=1 ms, V_th,tau_=-56 mV, V_slope,tau_=15 mV; in the Alpha/Beta model, p=1, k_α_=0.32 ms^-1^, V_α_=-52 mV, S_α_=-4 mV, k_β_=-0.28, V_β_ =-25 mV, S_β_=-5 mV. For a given G*_Kir,max_* value, the Hodgkin-Huxley Tau and Alpha/Beta models produced nearly identical results, therefore the data were pooled.

#### Analysis of membrane properties

The input resistance was estimated via the relation between the voltage response to the -50 to -10 pA current steps and the current step amplitude, which was well-fit by a linear relation in each PN. The membrane time constant was estimated using single exponential functions fit to the averaged voltage response to hyperpolarizing current steps of -30 to -10 pA in amplitude. This measure approximates the actual membrane time constant, which, in a passive neuron with complex geometry, is the time constant of the slowest component of a multiexponential time course (Spruston et al., 1994). The current threshold, or rheobase, was defined as the first current step amplitude producing at least one spike in both repeats for that current level. The voltage threshold to fire an action potential (AP threshold) was defined based on derivatives of the membrane potential (Henze et al., 2000). The afterhyperpolarization (AHP) amplitude was defined as the voltage from AP threshold to the trough of the AHP. The AP duration at half maximal amplitude (AP halfwidth) was defined as the duration of the AP at 50% of its maximal amplitude. All AP parameters were estimated from single APs (2 repeats per neuron) and measured on the first AP fired in response to rheobase current stimulation. The mean firing frequency and the spike probability were calculated only for experiments when 1 s current steps were used. The slope of the relation between mean firing frequency and current step amplitude (f–I plot) was estimated from the linear region of the plots of mean firing frequency versus stimulus amplitude, typically up to 100 pA above rheobase. The mean firing frequency was calculated from the number of APs evoked by the 1 s stimulus, averaged for the 2 repeats of each stimulus amplitude.

#### Voltage clamp

Recordings were obtained using Multiclamp 700A or 700B amplifiers (Axon Instruments) operating in voltage clamp mode. In both voltage and current clamp recordings the data were digitized at 50 kHz with Power 1401 digital-to-analog converters (Cambridge Electronic Design, CED, UK), using Signal 5 or Signal 7 software (CED). Recordings were performed without employing series resistance (R_series_) compensation, while the R_series_ was measured offline using inhouse written Signal scripts (Miyamae et al., 2017). In the experiments recording sIPSC recordings, the initial R_series_ was: dT-cells, 10.1±5.4 MΩ; dT+ cells, 9.9±4.1 MΩ (p=0.862, Mann-Whitney U test). In sEPSC recordings the initial R_series_ values was: dT-cells, 12.4±4.4 MΩ; dT+ cells, 14.0±5.9 MΩ, (F_(1,17.4)_=0.028, p=0.284, Linear Mixed Model). The R_series_ was monitored throughout recordings using small depolarizing or hyperpolarizing voltage steps (10 mV, 50 ms) delivered near the onset of each sweep. Sweeps were accepted for analysis only if the R_series_ increased less than 20% of the initial value. Synaptic currents were recorded holding the somatic membrane potential at -80 mV. Using NeuroMatic software (Rothman and Silver, 2018), with a sliding threshold search algorithm (Kudoh and Taguchi, 2002), we detected ∼200 synaptic current events per PN, measuring: 1) event frequency (number of events detected divided by the time window analyzed; 2) the average peak amplitude and 3) the individual sIPSC or sEPSC amplitudes, used to build the histograms of distribution.

### Histological processing and morphological reconstruction of biocytin-filled neurons

The p-formaldehyde-fixed slices containing the biocytin-filled neurons were stored at −80 °C in a cryo-protection solution (33% glycerol, 33% ethylene glycol, in 0.1 M PBS) until processed to visualize the biocytin. cells. For biocytin visualization, the slices were resectioned at 60 μm, incubated with 1% H_2_O_2_, and immersed in blocking serum containing 0.5% Triton X-100 for 2–3 h at room temperature. The tissue was then rinsed and incubated with the avidin–biotin– peroxidase complex (1:100; Vector Laboratories) in PBS for 4 h at room temperature. Sections were rinsed, stained with the Nickel-enhanced 3,3′-diaminobenzidine chromogen, mounted on gelatin-coated glass slides, dehydrated, and cover slipped. Three-dimensional reconstructions of the dendritic arbor were performed using the Neurolucida tracing system (MBF Bioscience). Dendritic spines were identified using differential interference contrast images of the biocytin-filled dendrites (González-Burgos et al., 2019; Gonzalez-Burgos et al., 2023). For the measurements of spine density, a single primary basal dendrite was randomly selected for each PN, and spines were reconstructed throughout the entire length of that dendrite. The peak spine density was defined as the highest spine number per micron value observed in each basal dendrite analyzed. The mean spine density was the average of the values of spine number per micron obtained for each basal dendrite analyzed.

### Statistical analysis

The data were expressed as mean±SD except if otherwise indicated. To assess normality of the distribution of the data we used the Shapiro-Wilk tests. If normality of distribution was rejected, the Shapiro-Wilk tests were repeated after applying natural logarithm (Ln) transformation of the data. Comparisons were performed with Linear Mixed Model or, when the data were obtained from a single animal, using Student’s t-test. If normality of the distribution was rejected after Ln transformation of the data, comparisons were performed using Generalized Linear Mixed Model, reporting the values of Chi Square Tests. When frequentist (p value-based) comparisons did not produce significant differences, we applied Bayesian tests as described elsewhere (Gonzalez-Burgos et al., 2023), using the Bayes Factor BF_10_, which quantifies the probability that the data are accounted for by either the null or the alternative hypothesis (Keysers et al., 2020). Statistical comparisons were performed using R code running under the JASP interface (https://jasp-stats.org/), except the Kolmogorov-Smirnov tests, performed using GraphPad Prism.

To obtain estimates of the probability density function of the distributions of sIPSC and sEPSC amplitude values, we used kernel smoothing to apply kernel density estimation implemented using scripts in OriginPro (OriginLab Co., MA USA). The optimal kernel bandwidth parameter ω was obtained for each data set by best fit of the kernelwidth function described by equation 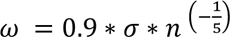, where n is the size of the dataset vector and σ = min (σ_s_, IQR/1.349) σ_s_ being the standard deviation of the dataset and IQR the interquartile range of the dataset. The kernel width ω obtained for each data set was used to compute the probability density function using equation 2: 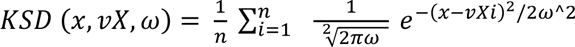, where n is the size of the dataset vector vX, ω is the kernel bandwidth and vXi is the ith element in the dataset vector vX. Probability density function fits were obtained for the distributions of sIPSC and sEPSC amplitude in each PN and averaged across cells to build the graphs in Figures 2D and 3D.

## Results

### Effects of Kir2.1 overexpression on DLPFC PN excitability

To decrease neuronal excitability, we overexpressed Kir2.1 in area 46 of monkey DLPFC by microinjecting an AAV which constitutively-expressed Cre recombinase and GFP under the CAMK2A promoter and another AAV encoding Cre-mediated expression of Kir2.1 and the fluorescent reporter dTomato (Fig 1A). This study is the first to use AAV-driven ion channel gene expression *in vivo* to manipulate the electrophysiology of primate neocortex neurons. Because we used the Kir2.1-encoding DNA sequence (KCNJ2) from the mouse genome (Xue et al., 2014), first we confirmed AAV-mediated expression of KCNJ2 in monkey DLPFC. In frozen DLPFC sections near and around AAV microinjections, we used fluorescent in situ hybridization (FISH), with probes for KCNJ2 and SLC17A7 (which encodes the PN specific marker VGluT1) mRNAs, to identify PNs with Kir2.1 expression. In tissue sections obtained near AAV injection sites, at 7-12 weeks post AAV microinjections, many neurons labeled by the SLC17A7 probe also were labeled for KCNJ2 (Extended data Fig 2A), whereas in tissue more distant from an injection site, KCNJ2-labeled cells were a smaller fraction of the cells labeled with the SLC17A7mRNA probe (Extended data Fig 2B), or were absent (Extended data Fig 2C). Thus, these data using FISH confirm that the two-AAV system employed produced robust expression of recombinant KCNJ2 mRNA in PNs.

**Figure 1.**
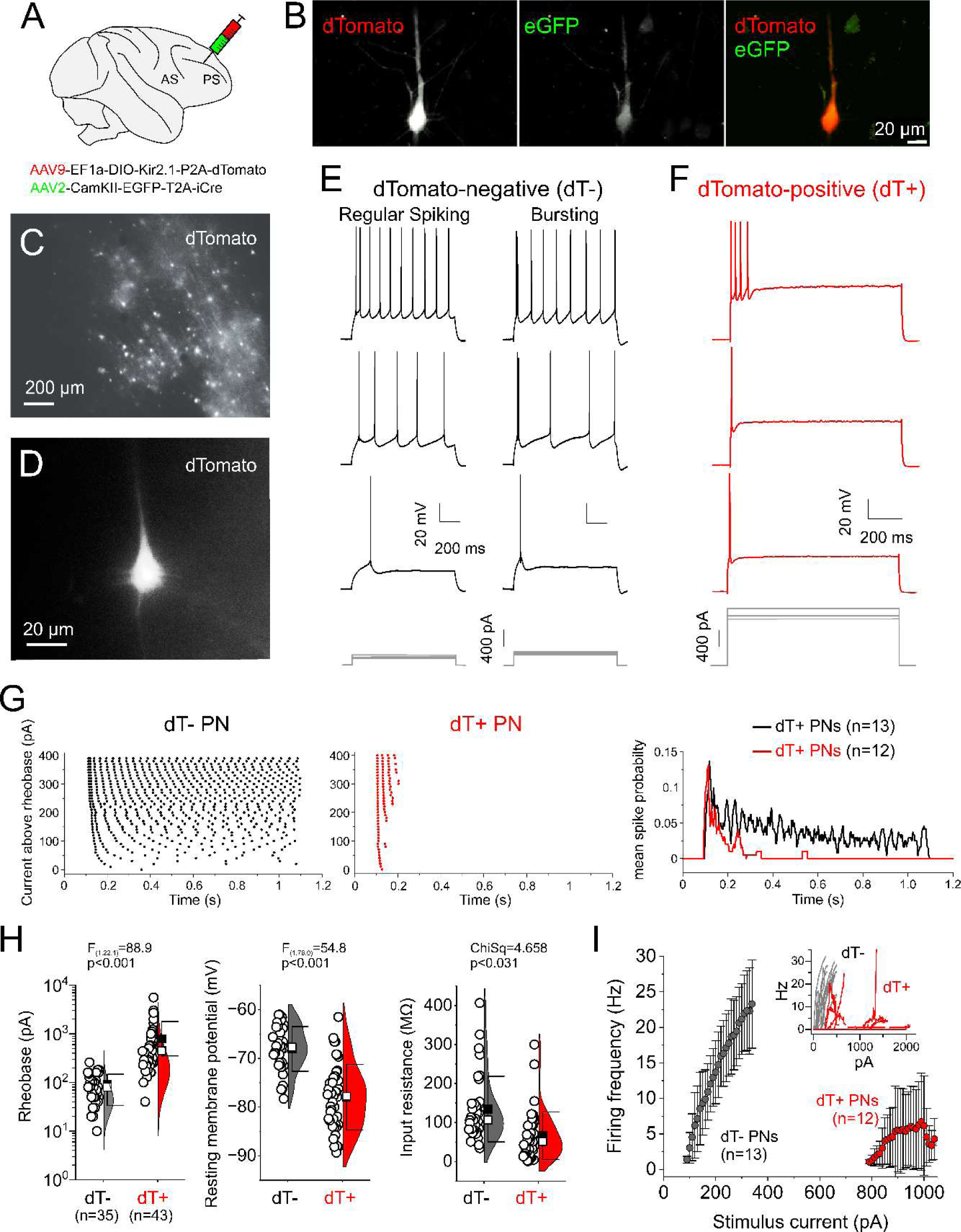
Kir2.1 overexpression affects the excitability and spiking pattern of PNs in DLPFC. **A**, Diagram indicating the two-AAV approach used to produce Cre recombinase-mediated Kir2.1 overexpression in DLPFC area 46. **B**, Confocal microscopy images of a pyramidal neuron (PN) expressing dTomato and GFP. **C**, Low magnification image of a field in an acute *ex* vivo DLPFC slice, showing a cluster of dTomato-positive (dT+) neurons. **D**, Higher magnification image of a dT+ PN visualized for patch clamp recording. **E**, Firing responses of two examples of dTomato-negative (dT-) PNs, showing typical Regular spiking and Burst spiking properties. **F**, Firing response of a dT+ PN with high rheobase current. **G**, Left and middle panels, raster plots illustrating the spiking patterns of dT+ and dT-PNs at different stimulus currents above rheobase. Right panel, spike probability throughout the duration of the 1 second stimulus averaged across dT- and dT+ PNs with stimulation current 250 pA above rheobase. **H**, Violin plots showing the differences in Rheobase (left), Resting membrane potential (middle) and Input resistance (right) between dT- and dT+ PNs. Data from experiments using 0.5 and 1 s-long current steps were pooled. **I**, Firing frequency versus stimulus current (f_I) plots calculated for dT- and dT+ PNs, showing data only from experiments using 1 s-long current steps. The inset shows the f_I plots for individual PNs, and the main graph depicts the average f_I plots, aligned to the mean rheobase of each group (dT-PNs: 90.5 pA; dT+ PNs: 788 pA).

**Figure 2.**
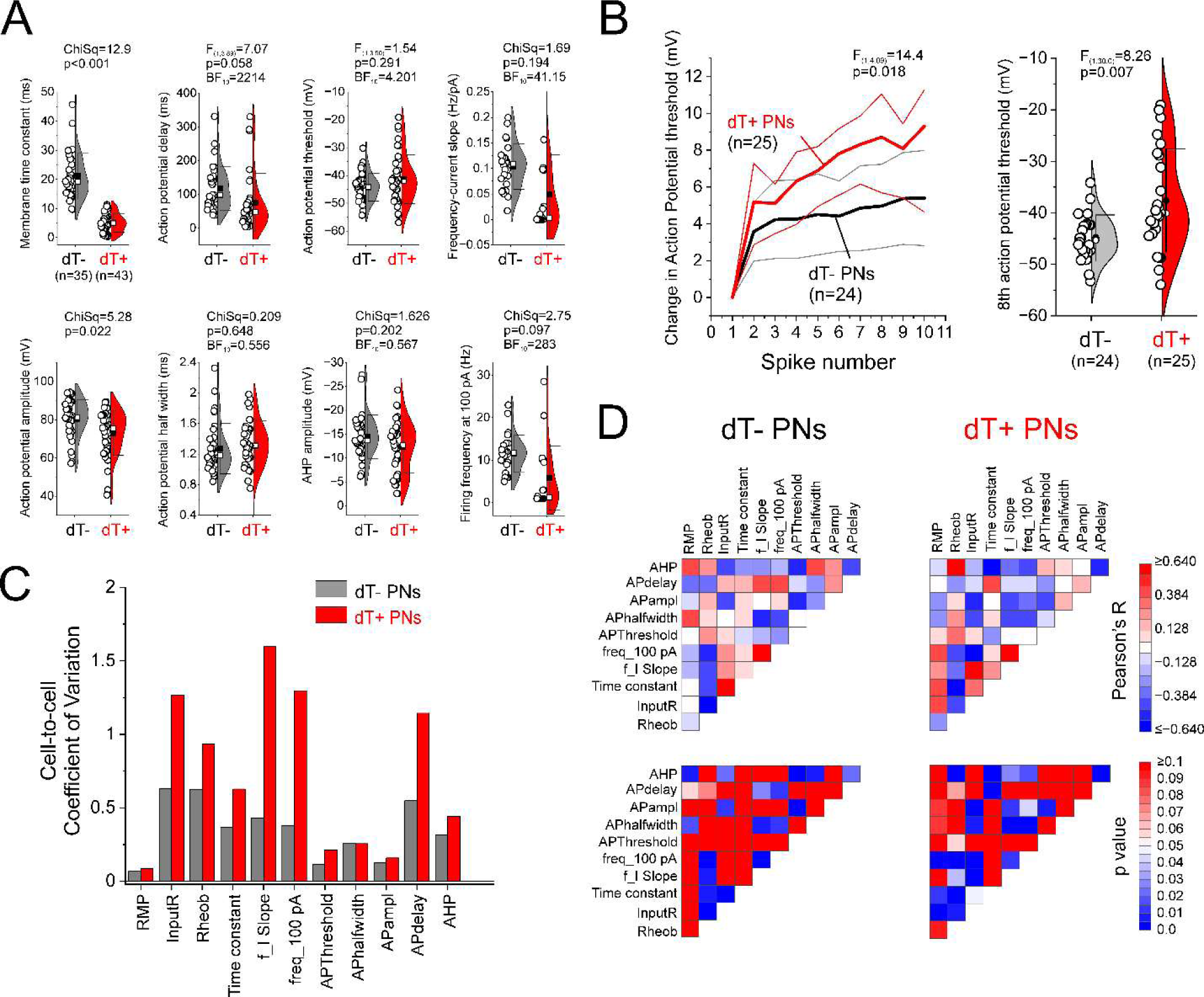
Membrane properties of dT- and dT+ PNs in DLPFC slices. **A**, Violin plots illustrating the differences in multiple membrane properties of dT- and dT+ PNs. **B**, Left panel, changes in the action potential threshold in dT- and dT+ PNs during spike trains, calculated subtracting the threshold of the first spike in each train. Right panel, violin plots depicting the AP threshold for the 8^th^ spike in the trains in dT- and dT+ PNs. **C**, The coefficient of variation between cells of the membrane properties of dT- and dT+ PNs. **D**, Pearson’s correlation coefficients (top heatmaps) and corresponding correlation p values (bottom heatmaps) estimated for the electrophysiological membrane properties of dT- (left column) and dT+ (right column) DLPFC PNs.

**Figure 3.**
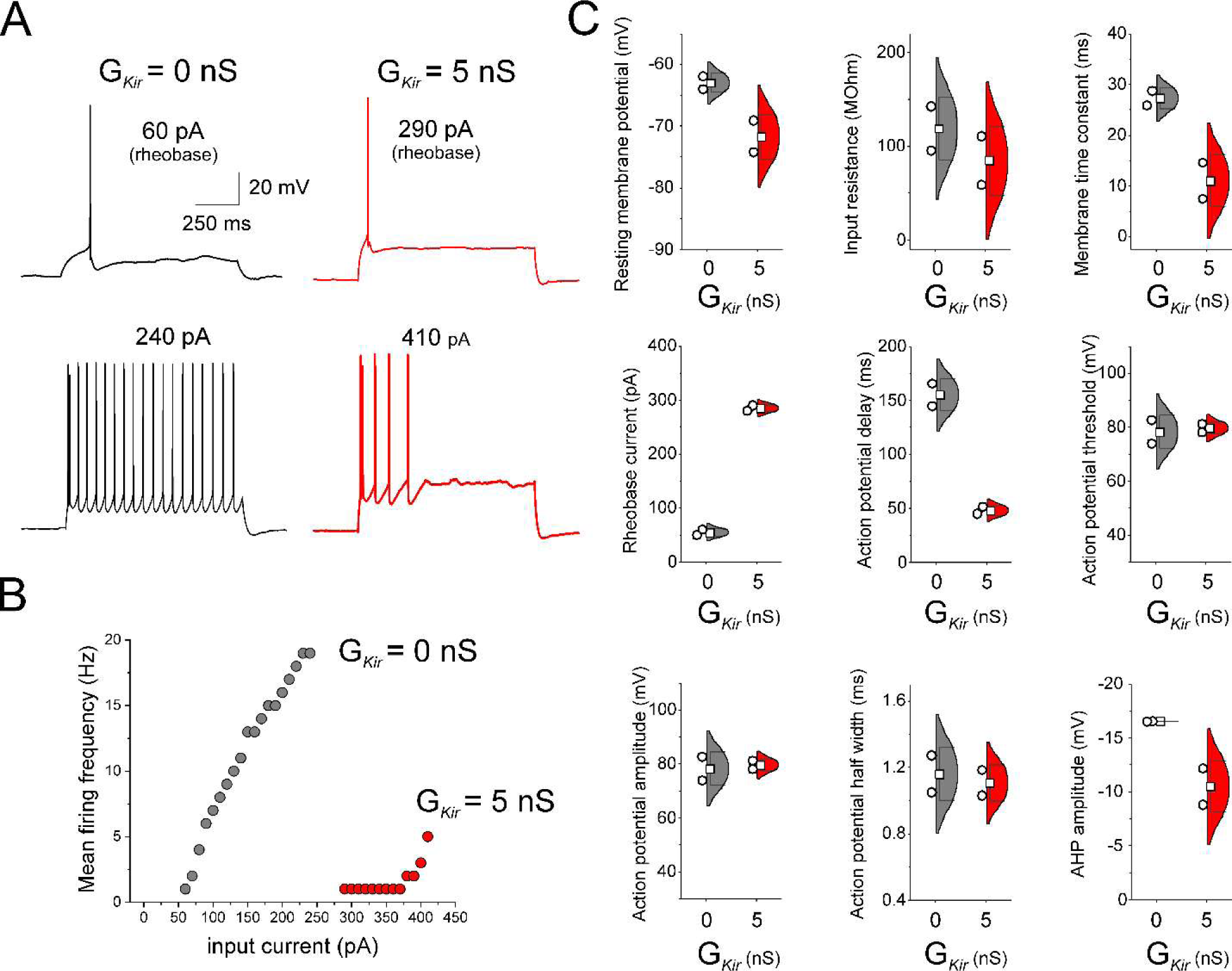
Dynamic clamp addition of Kir conductance (G*_Kir_*) to DLPFC PNs closely mimicked the membrane properties of dT+ PNs. **A,** Examples of recordings from PNs of monkey DLPFC before (G*_Kir_* = 0 nS) and during (G*_Kir_* = 5 nS) the addition of G*_kir_*. **B**, Input current versus firing frequency curve for a PN from monkey DLPFC before and after the addition of G*_Kir_*. **C**, Violin plots of membrane properties show that in a small sample of PNs from monkey DLPFC addition of G*_kir_* closely mimicked the membrane properties of dT+ PNs.

At 7-12 weeks post AAV microinjections, PNs expressing the GFP and dTomato fluorescent proteins were detectable in fixed tissue sections (Fig 1B), whereas in both acute living slices (Fig 1C) and fixed sections (Extended data Fig 3) adjacent to AAV injection sites, we observed clusters of dTomato-expressing (dT+) neurons. To obtain patch clamp recordings, single dT+ PNs were visualized and targeted in the living brain slices from DLPFC area 46 (Fig 1D). To confirm that Kir2.1 overexpression reduced the excitability of DLPFC PNs, we characterized the membrane properties of the dT+ PNs using current clamp recordings. In the acute brain slices, neither dT+ PNs nor PNs with undetectable dTomato fluorescence (dT-) showed spontaneous firing activity in the absence of stimulation. With depolarizing current injection, the dT-PNs showed firing responses (Fig 1E) very similar to those in the DLPFC of experimentally-naïve monkeys (Chang and Luebke, 2007; Zaitsev et al., 2012; González-Burgos et al., 2019; Medalla et al., 2022). In contrast, dT+ PNs required much stronger stimulation to produce action potentials (APs) and showed significantly altered firing responses (Fig 1F), often failing to fire persistently through the stimulus duration (Fig 1F,G). The dT+ PNs had higher rheobase, more negative resting potential and lower input resistance (Fig 1H), which likely contribute to their strongly reduced AP output (Fig 1I). The dT+ PNs displayed additional alterations in membrane properties (Fig 2), including a progressive increase in AP threshold throughout spike trains (Fig 2B), which may contribute to their failure to fire persistently (Fig 1F,G). Most membrane properties had higher cell-to-cell coefficient of variation in dT+ neurons (Fig 2C), consistent with variable Kir2.1 overexpression levels across PNs. Interestingly, as in human neocortex (Planert et al., 2023), in dT-PNs multiple membrane properties were significantly correlated, and many of these correlations were substantially altered in dT+ PNs (Fig 2D). Hence, Kir2.1 overexpression markedly reduced the excitability and globally altered the electrophysiology of DLPFC PNs.

The electrophysiological properties of dT+ PNs could be due exclusively to the biophysical effects of overexpressing Kir2.1 channels in the plasma membrane, or could reflect, in addition to the biophysical effects of Kir2.1 channels, alterations in other ion channels caused by the long-term (7-12 weeks) decrease of PN excitability *in vivo*. To characterize the biophysical effects of overexpressing Kir2.1 channels, we introduced Kir conductance (G*_Kir_*) using dynamic clamp (Amarillo et al., 2018; Szegedi et al., 2023) in dT-PNs from monkey DLPFC and mouse PFC. We found that adding G*_Kir_* reduced PN excitability while very closely mimicking the alterations in membrane properties in dT+ PNs (Fig 3,4), and in a G*_Kir_* dose-dependent manner (Fig 4C,D). Moreover, most PN membrane properties had substantially higher CV when variable amounts (1-20 nS) instead of a constant quantity (10 nS) of G*_Kir_* were introduced across neurons (Fig 4E). These data support the idea that the substantial cell-to-cell variability of membrane properties in dT+ neurons (Fig 2C) is caused by variable Kir2.1 overexpression levels. The similar properties of dT+ PNs and PNs with G*_Kir_* added by dynamic clamp show that our AAV-mediated silencing approach decreased PN excitability largely by highly specific effects of the AAV-encoded Kir2.1 channels. Moreover, these data suggest that 7-12 week-long overexpression *in vivo* did not cause major biophysical alterations beyond those produced by the presence of Kir2.1 channels in the plasma membrane.

**Figure 4.**
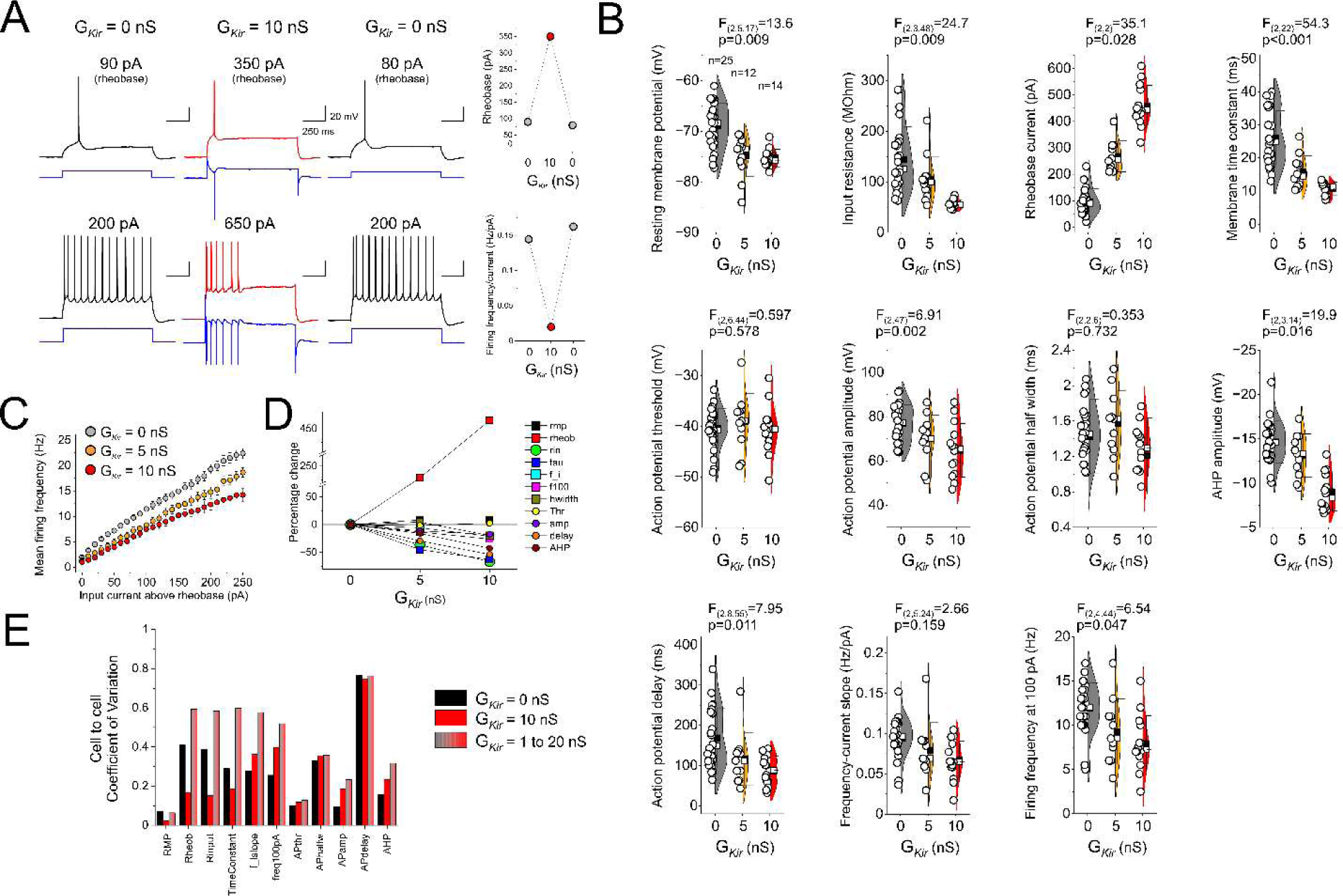
Dynamic clamp addition of Kir conductance (G*_Kir_*) to PNs from mouse PFC closely mimicked the membrane properties of dT+ PNs. **A**, Examples of recordings from PNs of mouse PFC before (G*_Kir_* = 0 nS) during (G*_Kir_* = 10 nS) and after (G*_Kir_* = 0 nS) the addition of G*_Kir_*. The plots to the right show that G*_Kir_* addition reversibly increased the Rheobase and decreased the firing rate per unit of current. **B**, Violin plots of membrane properties show that in a sample of PNs from mouse PFC addition of G*_Kir_* closely mimicked the membrane properties of dT+ PNs. **C**, Input current versus firing frequency curve for a PN from monkey DLPFC before and during the addition of G*_Kir_*. **D**, Addition of G*_Kir_* produced changes in membrane properties in a dose-dependent manner. **E**, The coefficient of variation of most membrane properties decreased with addition of a constant amount of G*_Kir_* (10 nS) but increased when variable G*_Kir_* levels (1-20 nS) were introduced with dynamic clamp.

### Effects of Kir2.1 overexpression on inhibitory and excitatory synapse strength

To determine if Kir2.1 overexpression affected synaptic inhibition in monkey DLPFC PNs, we recorded GABA_A_R-mediated spontaneous inhibitory synaptic currents (sIPSCs) (Fig 5A). Neither sIPSC frequency, mean sIPSC amplitude nor sIPSC amplitude distributions (Fig 5B-D) differed between dT+ and dT-PNs, suggesting that Kir2.1 overexpression did not affect inhibitory synaptic transmission. Albeit these data suggest an absence of difference in inhibitory synapse strength, it is possible that Kir2.1 overexpression produced changes in synaptic inhibition that were proportional to the magnitude of the reduction of excitability by Kir2.1 in each PN. Whereas detecting such a proportional change would ideally require measuring synaptic strength in each PN before and after Kir2.1 overexpression, such a proportional change might produce a correlation, in dT+ PNs, between parameters assessing excitability and GABA_A_ synaptic strength. However, scatter plots of rheobase, a strong index of Kir2.1-reduced excitability (Fig 1H, Fig 3C, Fig 4B), versus sIPSC amplitude or frequency did not reveal a consistent trend (Fig 5E,F), and the rheobase was not significantly correlated with sIPSC amplitude or frequency. Consistent with these findings, mean protein levels of the 67 kD isoform of glutamic acid decarboxylase (GAD67), the enzyme responsible for most GABA synthesis in the cortex, in axon terminals (as measured by fluorescent immunohistochemistry (Rocco et al., 2016)) did not differ (p = 0.64) between DLPFC tissue zones with (311±109 a.u.) or without (335 ± 107 a.u.) dT+ PNs.

**Figure 5.**
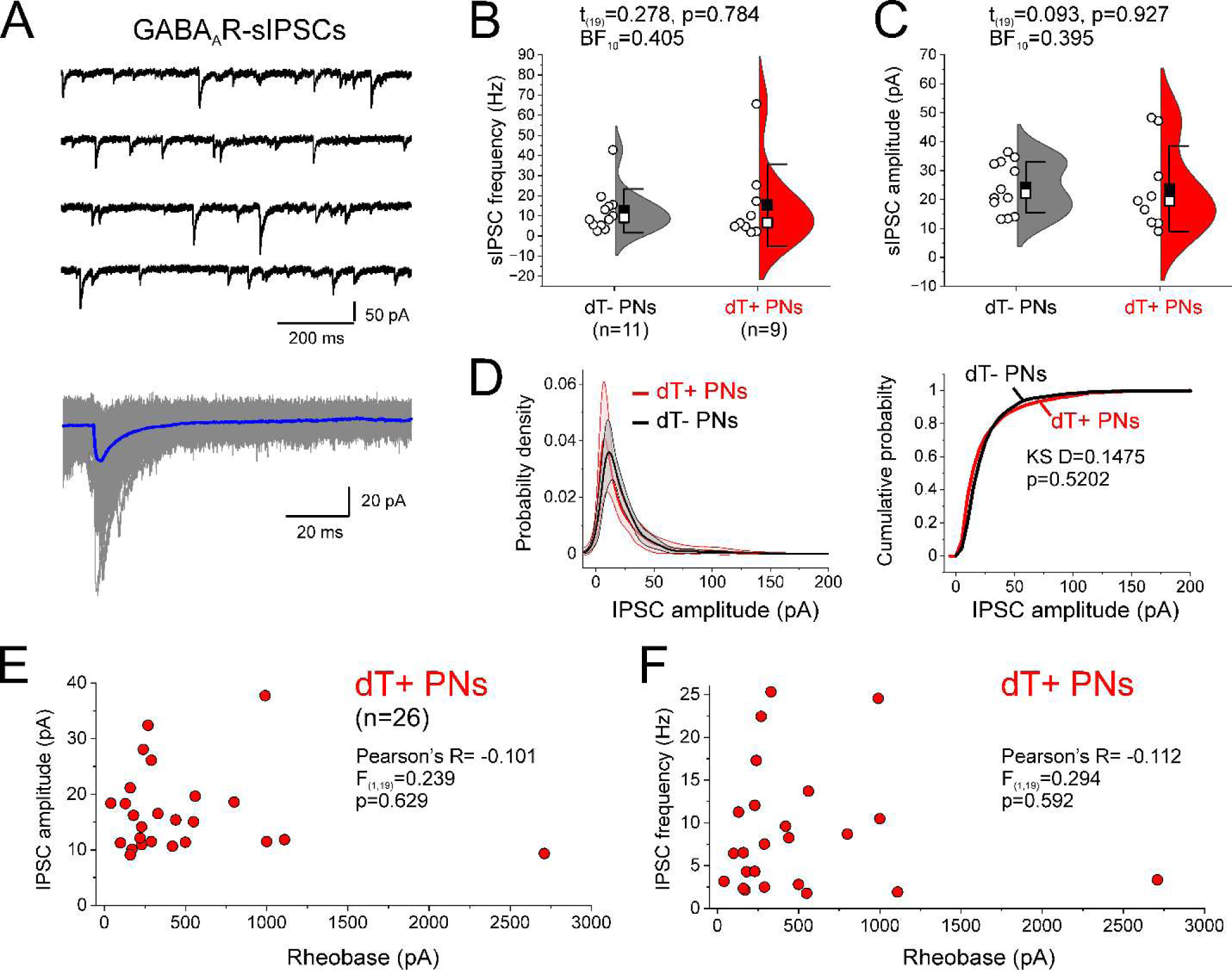
Kir2.1 overexpression did not affect the properties of GABA_A_R-mediated spontaneous IPSCs (sIPSCs) in DLPFC PNs. **A**, Top panel, example traces of the recorded sIPSCs. Bottom panel, multiple individual sIPSCs (gray traces) are shown superimposed together with their average waveform (blue trace). **B** and **C**, Violin plots comparing the sIPSC frequency (B) and sIPSC mean amplitude (C) in dT- and dT+ PNs. **D**, Left panel, histograms of distribution of sIPSC amplitudes in dT- and dT+ PNs. Shown are the mean±confidence intervals from probability density functions fit to the distribution of each PN. Right panel, cumulative probability distribution graphs for sIPSC amplitude. **E**, and **F**, Plots of sIPSC amplitude (E) and sIPSC frequency (F) versus rheobase measured in single dT+ DLPFC PNs. sIPSC amplitude and frequency data are from the same PNs.

To test whether Kir2.1 overexpression affected excitatory synaptic function we compared spontaneous excitatory synaptic currents (sEPSCs) recorded from dT+ and dT-PNs (Fig 6A). Whereas sEPSC frequency did not differ between dT+ and dT-PNs (Fig 6B), the mean sEPSC amplitude was significantly smaller in dT+ PNs (Fig 6C). Albeit the difference in mean sEPSC amplitude was relatively modest (Fig 6C), the distribution of sEPSC amplitudes showed a robust leftward shift in dT+ PNs (Fig 6D), suggesting that Kir2.1 overexpression decreased strength across the population of excitatory synapses. If the magnitude of the decrease in excitatory synaptic strength depended on the degree by which the postsynaptic excitability is reduced by Kir2.1 overexpression, then in dT+ PNs the sEPSC amplitude might be inversely correlated with the rheobase. We tested the correlation between the rheobase and the sEPSC amplitude and the most frequent sEPSC amplitude in each PN, given that the distributions of sEPSC amplitudes were markedly skewed (Fig 6D, Extended Data Fig 4), and thus the mean sEPSC amplitude does not represent the most frequent synapses, which might be the synapses most commonly driving PN activity. The most frequent sEPSC amplitude, estimated in each PN using probability density functions, and this parameter was significantly smaller (F_(1,21.6)_=6.620, p=0.018) in dT+ (5.9±1.7 pA, n=15) than dT- (7.9±3.0 pA, n=15) PNs. Neither the mean (Fig 6E) nor the most frequent (Fig 6F) sEPSC amplitudes were significantly correlated with the rheobase. Thus, the magnitude of the decrease in excitatory synaptic strength was not tightly associated with the degree of excitability reduction produced by *in* vivo Kir2.1 overexpression.

**Figure 6.**
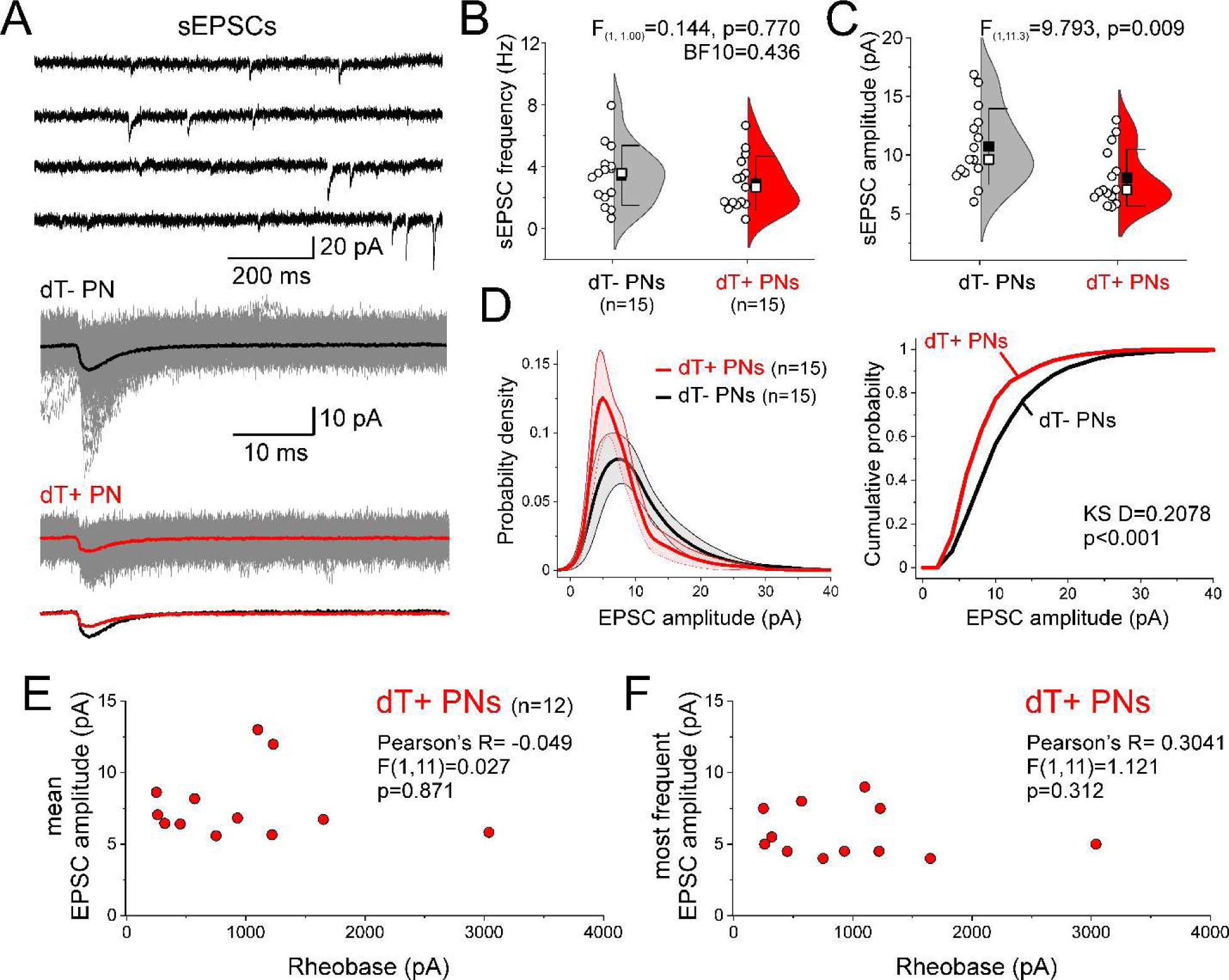
Kir2.1 overexpression increased the amplitude of spontaneous EPSCs (sEPSCs) in DLPFC PNs. **A**, Top panel, example traces of the recorded sEPSCs. Middle panel, multiple individual sEPSCs (gray traces) are shown superimposed together with their average waveforms for dT- (black trace) and dT+ (red trace) PNs. Bottom panel, the average waveforms are shown superimposed. **B** and **C**, Violin plots comparing the sEPSC frequency (B) and sEPSC mean amplitude (C) in dT- and dT+ PNs. **D**, Left panel, histograms of distribution of sEPSC amplitude in dT- and dT+ PNs. Shown are the mean±confidence intervals from probability density functions fit to the distribution of each PN. Right panel, cumulative probability distribution graphs for sEPSC amplitude. **E**, and **F**, Plots of mean sEPSC amplitude (E) and most frequent sEPSC amplitude (F) versus rheobase measured in single dT+ DLPFC PNs. The most frequent sEPSC amplitude was estimated using the probability density functions fit to the distribution of sEPSC amplitude in each PN.

Given that the distribution of the overexpressed Kir2.1 channels along the somato-dendritic plasma membrane of dT+ PNs is unknown, one possibility is that the lower amplitude of sEPSCs recorded from dT+ PNs using somatic voltage clamp was caused by greater space clamp errors due to dendritic Kir2.1 channel localization. Importantly, however, seminal studies employing computational modeling (Spruston et al., 1993) and experimental (Williams and Mitchell, 2008) approaches showed that decreasing the dendritic membrane resistance (as with Kir2.1 channels localized in dendrites), does not increase the space clamp errors during somatic recordings of distally-generated EPSCs. Hence, next we measured the rising phase slope of the sEPSCs, a kinetic parameter that depends on, among other factors, dendritic synapse location. Partly due to space clamp errors, rising speed is slower the more distal a synapse producing an EPSC is located (Spruston et al., 1993; Williams and Mitchell, 2008). We found that, as expected, the sEPSC rising phase slopes had wide distributions, but that the distributions were very similar in dT+ and dT-PNs (Extended Data Fig 5A). These results contradict the prediction that, if space clamp errors were larger in dT+ PNs, these cells would show a greater proportion of sEPSCs with slow rising phase. Moreover, contrary to the expectation that with worse space clamp errors, sEPSCs with slower rising phase slope would have a greater decrease in amplitude, the sEPSCs in dT+ PNs had smaller amplitude throughout the distribution of rising phase slopes (Extended Data Fig 5B).

To test whether the decrease in excitatory synaptic strength (Fig 6) was associated with changes in excitatory synapse quantities, we measured spine density in PN basal dendrites. Both dT+ and dT-PNs exhibited typical distributions (Elston and Rosa, 1997; Elston et al., 2011; Medalla and Luebke, 2015; González-Burgos et al., 2019) of dendritic spines across the basal dendrite length (Fig 7A,B), and exhibited typical mean and peak spine densities which did not differ between dT+ and dT-cells (Fig 7C,D). Moreover, both basal dendrite length and architecture were unaffected in dT+ PNs (Fig 7E-J).

**Figure 7.**
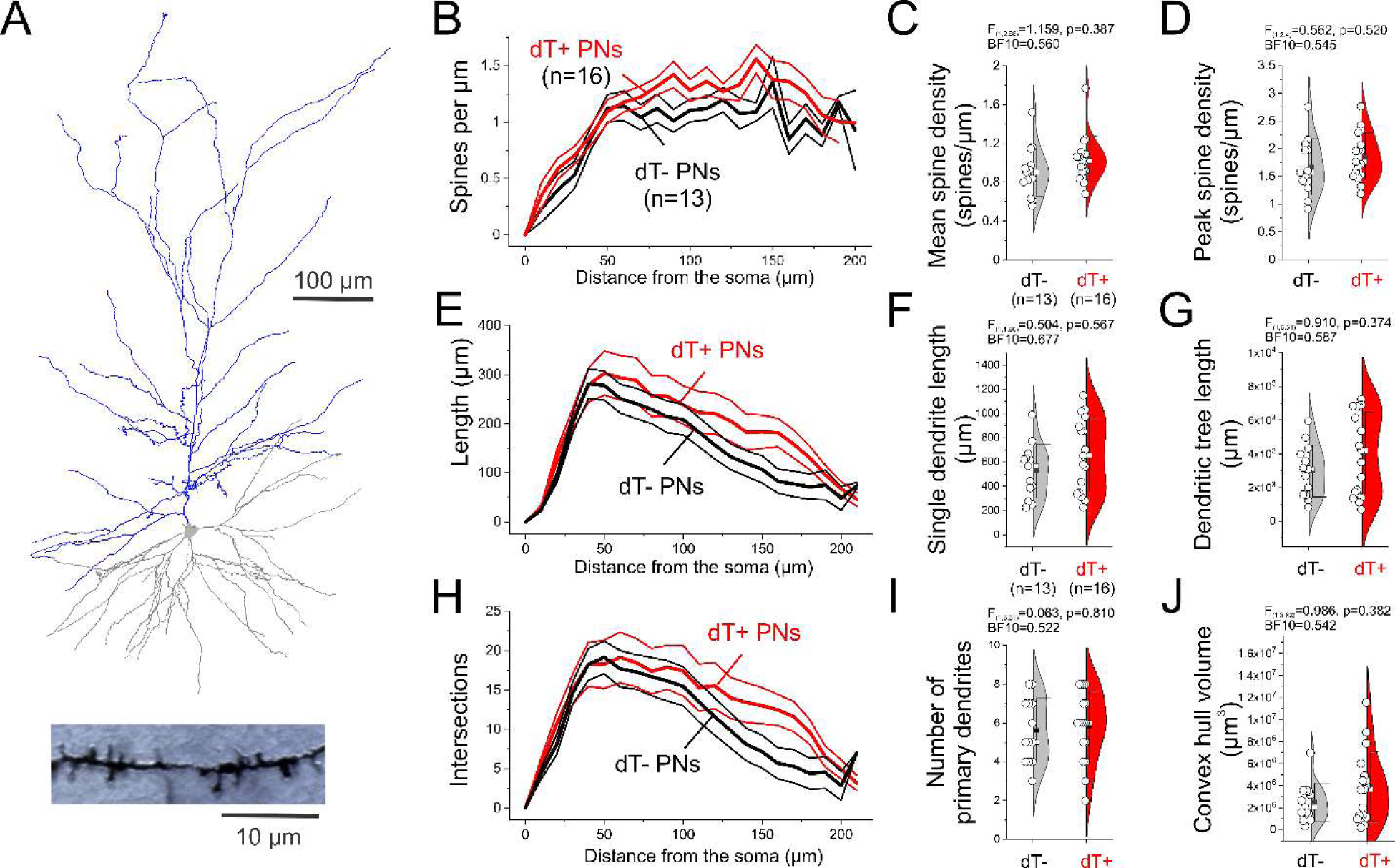
Kir2.1 overexpression did not affect dendritic tree properties in PNs from monkey DLPFC. **A**, Top panel, example of the reconstruction of the dendritic tree of a PN recorded in the DLPFC slices filled with biocytin during recording. Bottom panel, a short basal dendrite segment from a biocytin-filled PN, illustrating examples of dendritic spines. **B,** Plots of spine density as a function of distance from the soma. **C** and **D**, Violin plots illustrating mean and peak spine density, respectively. **E**, Plots of basal dendrite length as a function of distance from the soma. **F**, Violin plots of length of single basal dendrite. **G**, Violin plots of the total length of basal dendrites. **H**, Plots of dendritic tree intersections as a function of distance from the soma. **I,** Violin plots of number of primary basal dendrites. **J,** Violin plots of convex hull volume for the basal dendrites.

## Discussion

Data from previous studies suggest that, in schizophrenia, PNs from DLPFC have reduced activity. For instance, these neurons have a lower density of dendritic spines (Garey et al., 1998; Glantz and Lewis, 2000), the main site of excitatory drive onto PNs, and also reduced expression of activity-dependent genes (Weickert et al., 2003; Hashimoto et al., 2005) and of genes involved in energy production (Arion et al., 2015; Arion et al., 2017; Glausier et al., 2020). Reduced PN activity is consistent with the reduced activation and lower gamma oscillation power during cognitive tasks in the DLPFC of people with schizophrenia (Cho et al., 2006; Minzenberg et al., 2009; Boudewyn et al., 2020; Smucny et al., 2023). These observations prompted us to test whether reducing the activity of PNs in the DLPFC affects the strength of inhibitory and excitatory synaptic inputs onto these neurons.

We found that Kir2.1 overexpression in macaque DLPFC *in vivo* decreased PN excitability *ex vivo*, but without affecting inhibitory synaptic currents. Our results therefore suggest that reducing the excitability of a PN *in vivo* is not sufficient to cause changes in the inhibitory inputs onto that cell; these findings may be unique to the mature primate DLPFC as a similar Kir2.1-mediated experimental manipulation in early postnatal visual cortex from mice weakened inhibitory synapses (Xue et al., 2014). Our findings suggest that in schizophrenia other pathogenic processes (e.g., cell-autonomous disturbances in interneurons (Chung et al., 2016)) might be required, in addition to lower PN activity, to alter inhibitory synapses (Gonzalez-Burgos et al., 2015a). However, it is also possible that the 7-12 weeks of Kir2.1 overexpression in the present study were not sufficient to capture the cumulative effects on synaptic inhibition of the much longer period of PN hypoactivity likely present in schizophrenia (Smucny et al., 2022). Furthermore, the alterations of inhibitory synapses reported in schizophrenia might also be accompanied by lower interneuron activity (Smucny et al., 2022), but it is unclear if in the present study the percentage of PNs overexpressing Kir2.1 was sufficient to significantly reduce the recruitment of interneurons via excitatory-to-inhibitory synaptic connections. Thus, future studies assessing interneuron properties and/or involving Kir2.1 overexpression directly in interneurons might be informative. Interestingly, although inhibitory synapses are regulated via homeostatic mechanisms, in several studies manipulations producing excitatory synapse up-scaling did not change inhibitory synapse strength or changed this strength following a markedly different timing (Wen and Turrigiano, 2024). Moreover, it is possible that Kir2.1 overexpression changes the net effect of synaptic inhibition on DLPFC neurons without changing GABA synapse strength, but instead regulating interneuron excitability or the strength of the excitatory synapses recruiting the interneurons (Wen and Turrigiano, 2024). Finally, it is also possible that lower PN activity affects the inhibitory synapses from specific interneuron subtypes, and that such an effect is not detectable by recording spontaneous IPSCs.

Our findings that Kir2.1 overexpression in macaque DLPFC *in vivo* decreased excitatory synaptic strength *ex vivo* contrast with previous reports showing that experimentally reducing neuronal activity, for a shorter duration in the visual cortex of relatively young mice, up-regulated excitatory synapse strength (Xue et al., 2014; Wen and Turrigiano, 2021). Importantly, the absence of excitatory synapse upscaling in dT+ PNs seems to be at variance with the model of homeostatic regulation of excitatory synaptic strength (Wen and Turrigiano, 2024). The dissimilar findings suggest that the alterations of excitatory synaptic strength produced by lower PN activity may differ based on age, cortical area, species and/or the duration and magnitude of lower PN activity. One possibility is that Kir2.1 overexpression affected the mechanisms of spike timing-dependent plasticity in a manner that relatively favored depression over potentiation of synaptic strength. For instance, by markedly decreasing the total spike output of dT+ PNs, Kir2.1 overexpression could disrupt the patterns of paired/coincident pre and postsynaptic spikes that commonly maintain excitatory synapse strength.

Whereas multiple studies reported that network activity during cognitive tasks is reduced in the DLPFC in schizophrenia (Cho et al., 2006; Minzenberg et al., 2009; Boudewyn et al., 2020; Smucny et al., 2023), the extent to which single PN activity is reduced is unknown. Importantly, to our knowledge neither Kir2.1 channel protein levels nor KCNJ2 mRNA levels are affected in PNs in schizophrenia (Arion et al., 2015). However, if the schizophrenia disease effect reduces PN activity to levels comparable to the reduction produced by Kir2.1 overexpression in this study, then our findings suggest the novel prediction that PNs with lower activity in schizophrenia would also display reduced excitatory synapse strength. Therefore, the combined effects of lower synapse number and strength (which might occur sequentially during postnatal development if excessive synaptic pruning, resulting in lower spine density, is the proximal cause of PN hypoactivity) may lead to the emergence of DLPFC microcircuit dysfunction in the illness, and interventions promoting potentiation of the weakened excitatory synapses could improve the associated cognitive deficits.

## Acknowledgments

We thank Jacqueline Breter for excellent assistance during performance of the surgical procedures. Funding: NIH Grant MH043784 (D. A. L.), NIH Grant MH117089 (M. X.), NSF Grant Neuronex 2015276 (G. G. B.).

---

**Extended Data Figure 1.**
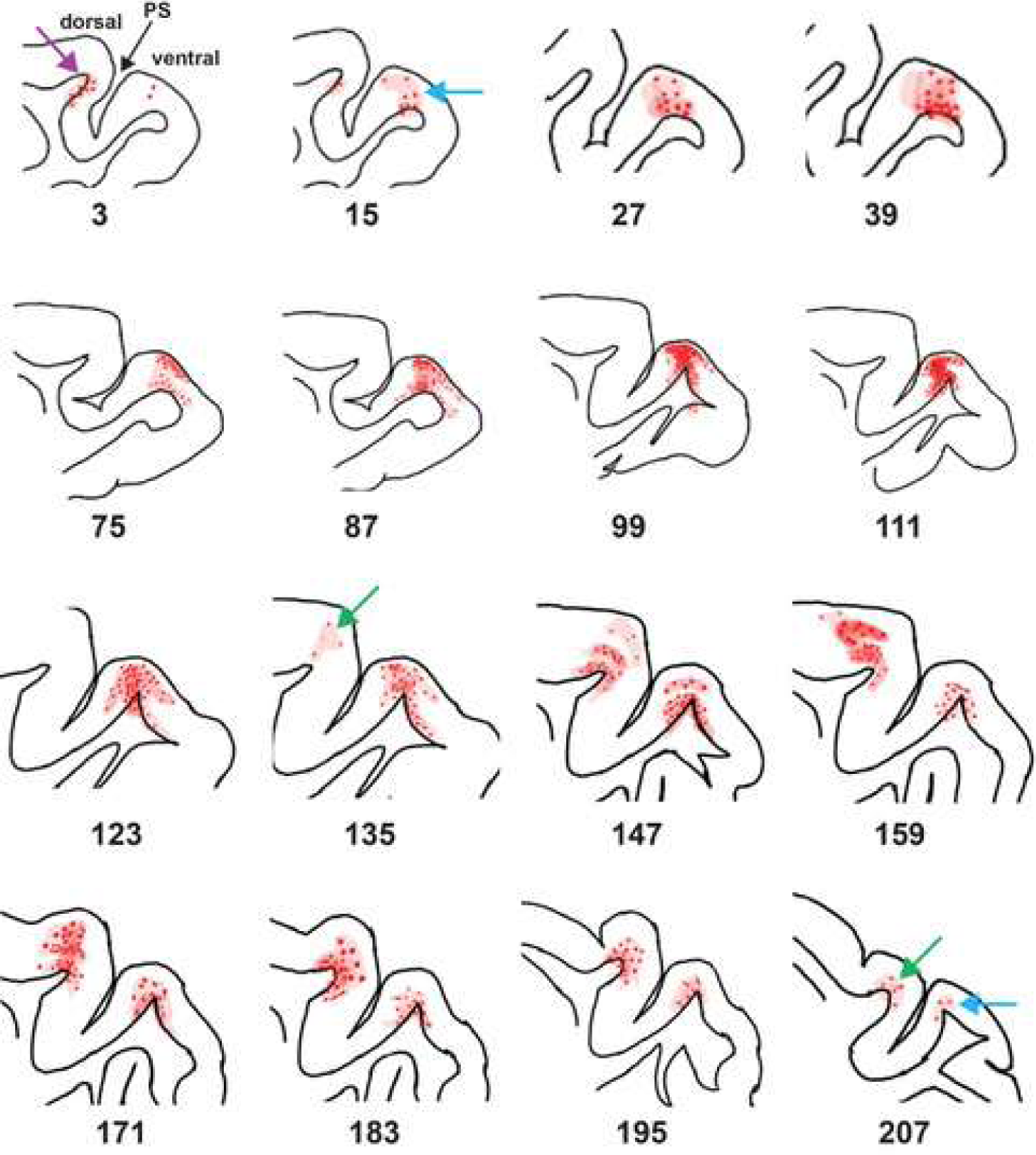
Schematic drawing illustrating the distribution of cells expressing dTomato (dT+ cells) in a representative AAV injection (subject vi002). Red shading represents general fluorescence, while red dots represent clusters of dT+ cells. Numbers under each schematic designate the section number from the tissue block, which was cut to exhaustion. In section 3, the end of an injection (purple arrow) can be seen in the dorsal bank of the principal sulcus (PS), while the beginning of another injection on the ventral bank, is marked with a blue arrow in section 15. The beginning of a third injection site is present on the dorsal bank of section 135 (green arrow).

**Extended Data Figure 2.**
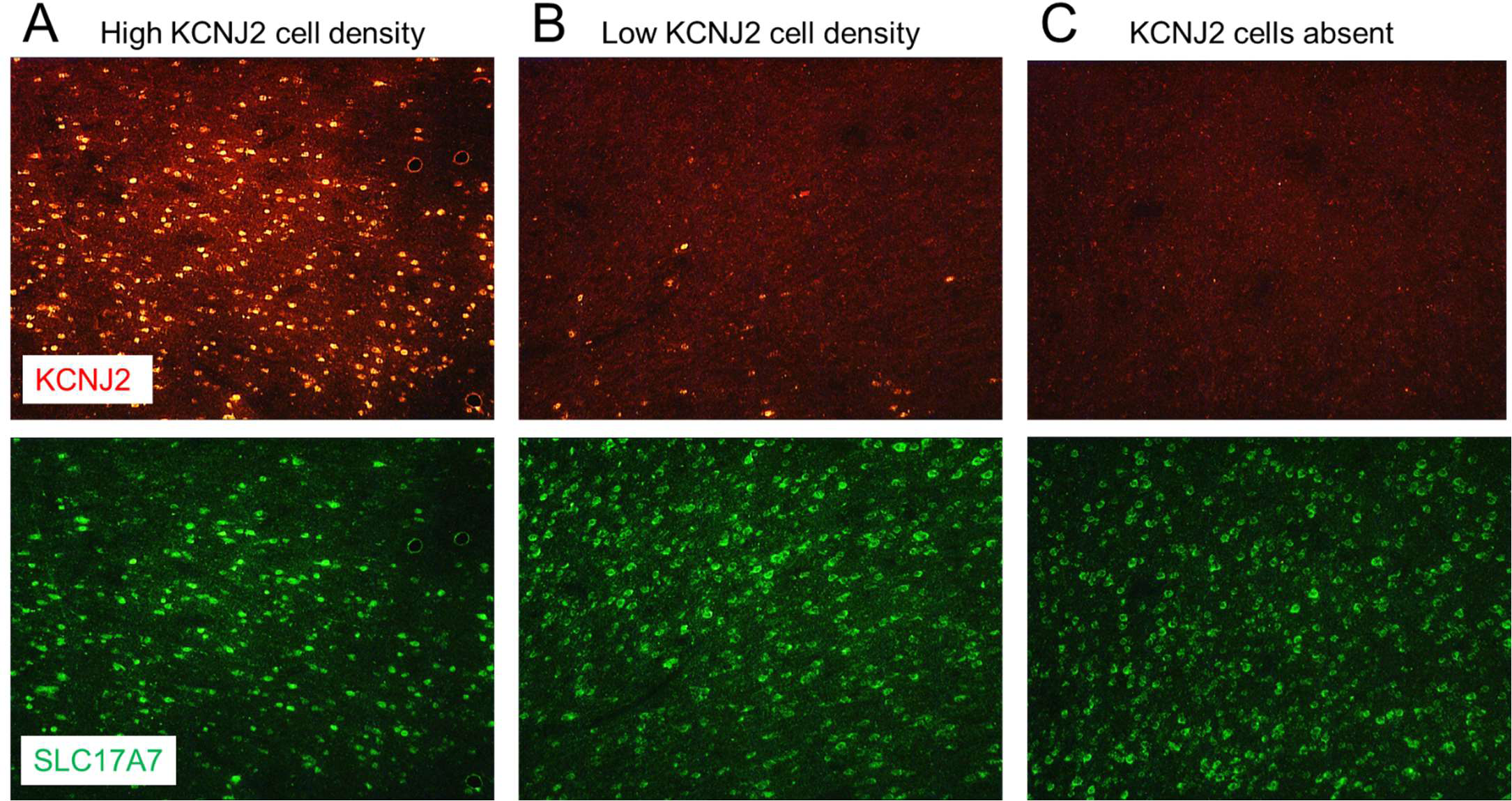
Fluorescence In Situ Hibridization (FISH) assessment of expression of recombinant KCNJ2 mRNA with AAV injections into DLPFC area 46. **A)** Example of neurons labeled in a tissue section near an AAV injection site, by fluorescent probes detecting KCNJ2 (red) and SLC17A7 (green), encoding Kir2.1 channels and the pyramidal neuron marker VGluT1, respectively. The images were captured from a tissue section near an AAV injection site, depicting high density of cells labeled by both probes. **B)** and **C)** identical analysis for tissue sections at intermediate (B) and far (C) distances from the same injection site.

**Extended Data Figure 3.**
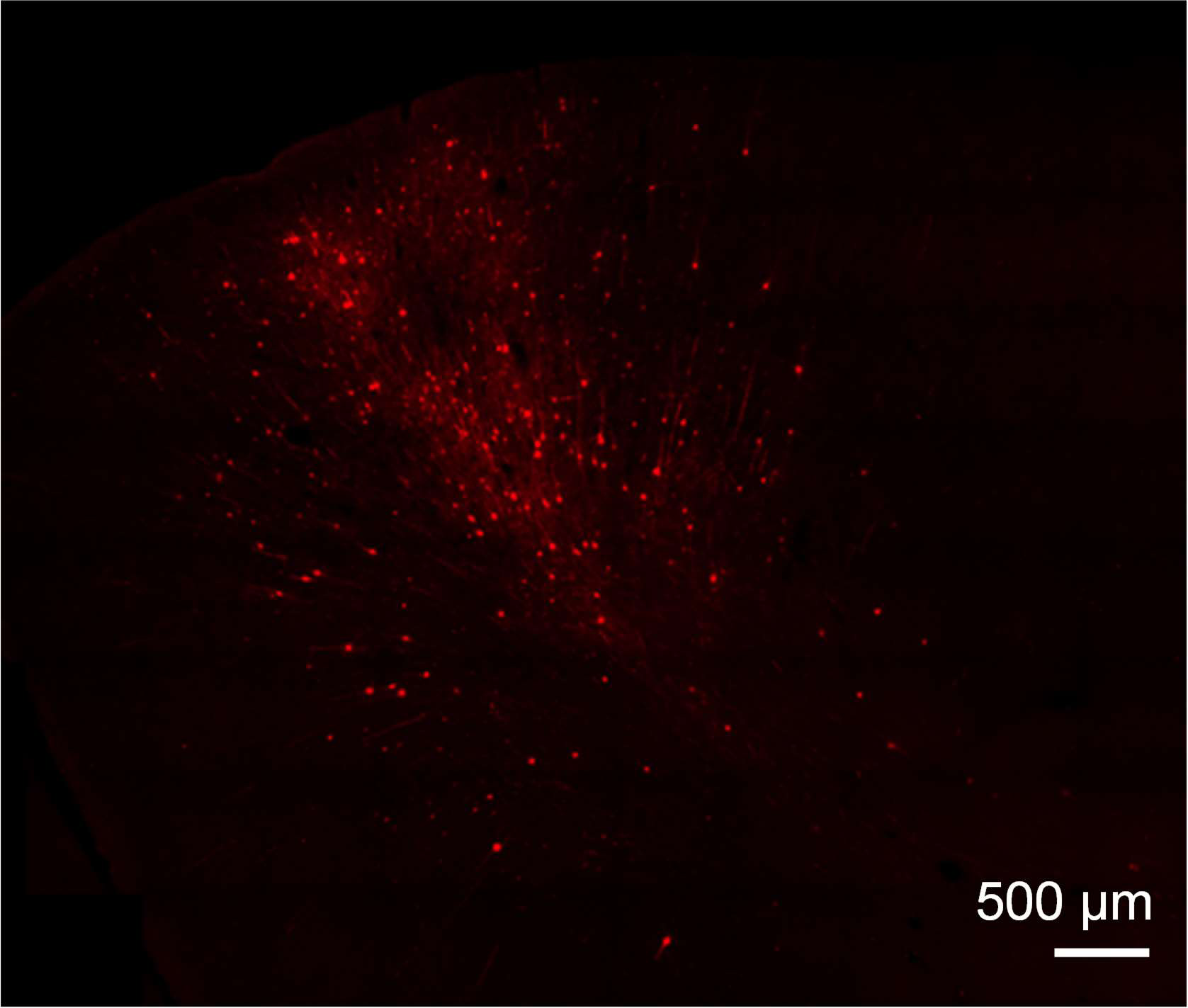
Low magnification image of a field in a formaldehyde-fixed section from DLPFC tissue adjacent to an AAV injection site, showing a cluster of dTomato-positive (dT+) neurons.

**Extended Data Figure 4.**
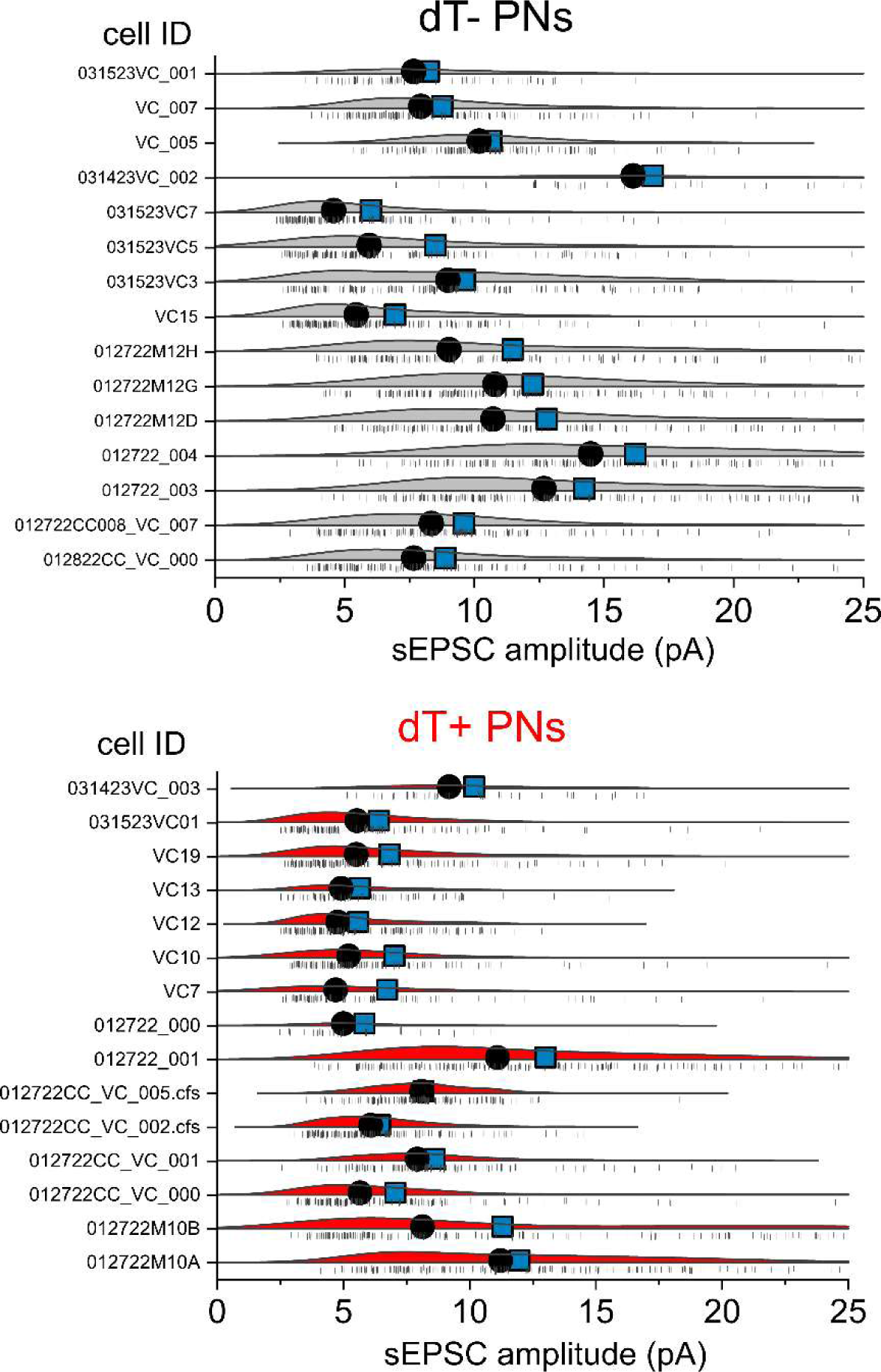
Distribution of sEPSC amplitudes in individual dT- (top panel) and dT+ PNs (bottom panel) from monkey DLPFC. Shown are probability density functions fit to the distribution of sEPSC amplitudes in each PN, together with the location of individual sEPSCs in each distribution. The black dots and blue squares depict, respectively, the median and mean of the distributions. Note that due to the skewed shape of most distributions, neither median nor mean represent the most frequent sEPSC amplitude in each neuron, which was calculated as maximum value in the probability density functions fit to the data (see text for additional details).

**Extended data Figure 5.**
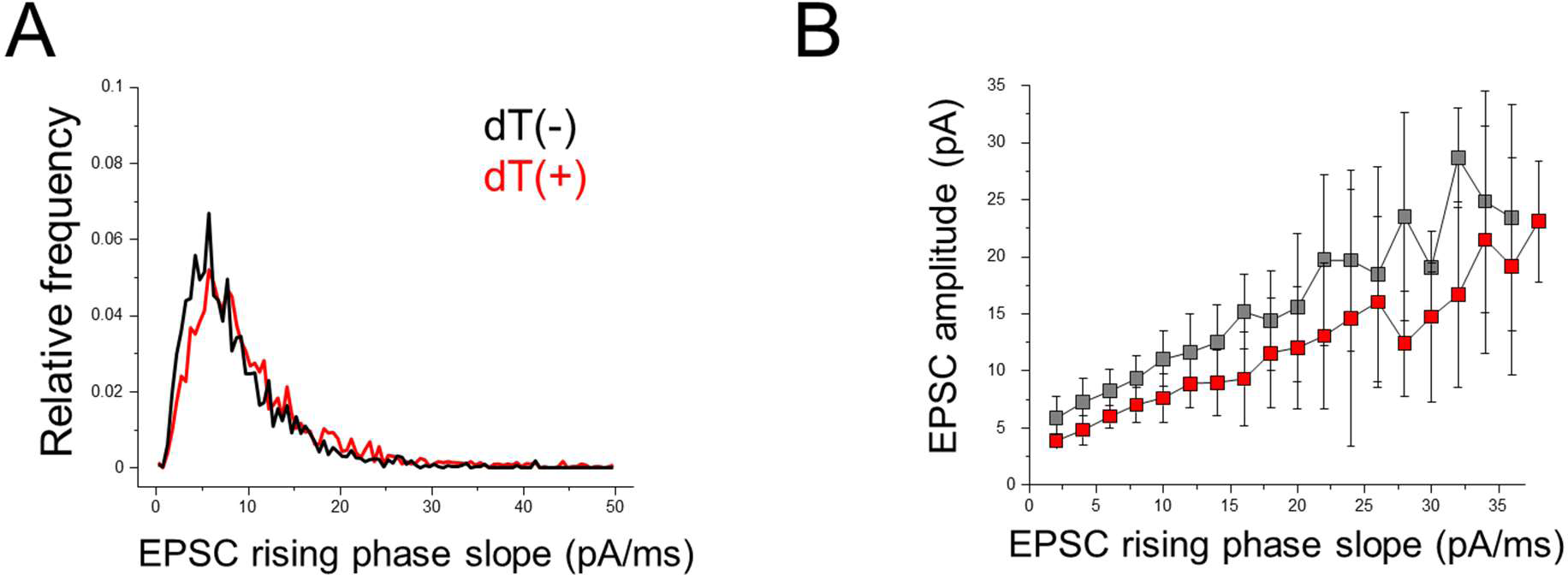
Analysis of the kinetics of rise of sEPSCs recorded from dT+ and dT-PNs. **A)** Distribution of sEPSC rise times in the entire data sample from recording sEPSCs in dT+ and dT-PNs. Note that the distribution of sEPSC rising phase slopes is inconsistent with slower rising phase, presumably due to worse space clamp error when voltage clamping dT+ PNs. **B)** Plots of sEPSC amplitude as a function of sEPSC rising phase slope, showing that the lower sEPSC amplitude in dT+ PNs, is not associated with slower rising phase values, as expected if the decrease in amplitude was due to worse space clamp error when voltage clamping dT+ PNs.

